# Large-Scale Search of Transcriptomic Read Sets with Sequence Bloom Trees

**DOI:** 10.1101/017087

**Authors:** Brad Solomon, Carleton Kingsford

**Affiliations:** Computational Biology Department, School of Computer Science, Carnegie Mellon University, Pittsburgh, PA 15213, USA

## Abstract

Enormous databases of short-read RNA-seq sequencing experiments such as the NIH Sequence Read Archive (SRA) are now available. However, these collections remain difficult to use due to the inability to search for a particular expressed sequence. A natural question is which of these experiments contain sequences that indicate the expression of a particular sequence such as a gene isoform, lncRNA, or uORF. However, at present this is a computationally demanding question at the scale of these databases.

We introduce an indexing scheme, the Sequence Bloom Tree (SBT), to support sequence-based querying of terabase-scale collections of thousands of short-read sequencing experiments. We apply SBT to the problem of finding conditions under which query transcripts are expressed. Our experiments are conducted on a set of 2652 publicly available RNA-seq experiments contained in the NIH for the breast, blood, and brain tissues, comprising 5 terabytes of sequence. SBTs of this size can be queried for a 1000 nt sequence in 19 minutes using less than 300 MB of RAM, over 100 times faster than standard usage of SRA-BLAST and 119 times faster than STAR. SBTs allow for fast identification of experiments with expressed novel isoforms, even if these isoforms were unknown at the time the SBT was built. We also provide some theoretical guidance about appropriate parameter selection in SBT and propose a sampling-based scheme for potentially scaling SBT to even larger collections of files. While SBT can handle any set of reads, we demonstrate the effectiveness of SBT by searching a large collection of blood, brain, and breast RNA-seq files for all 214,293 known human transcripts to identify tissue-specific transcripts.

The implementation used in the experiments below is in C++ and is available as open source at http://www.cs.cmu.edu/∼ckingsf/software/bloomtree.

## 1 Introduction

The amount of DNA and RNA short read sequence data that has been published world wide is huge. For example, the NIH’s sequence read archive (SRA) [14] alone contains 3,143,709,042,522,870 bases of sequence (about 3 petabases). This collection of experiments is a great resource for understanding genetic variation, and condition- and disease-specific gene function in ways the original depositors of the data did not anticipate. In aggregate too the data could provide the ability to answer questions that a single experiment could not answer. However, its use is hampered by its size and the inability to search experiments by sequence in meaningful ways. For example, a natural use of this data would exploit the 19,807 human RNA-seq short-read files in the SRA (representing individual sequencing runs) to understand the conditions under which a novel or hypothesized alternatively spliced isoform *q* is expressed by searching each experiment for enough reads matching *q* to indicate the *q* was present in the experiment. However, searching the entirety of such a database for expressed genes, novel isoforms, or homologous gene sequences has not been possible in reasonable computational time. This is the problem we tackle here.

We present and test a system to find the experiments in which a given query sequence *q* is likely expressed from among thousands of short-read sequencing experiments comprising terabases of sequence data. Such large-scale querying could be used to hypothesize functions for the thousands of long non-coding RNAs by identifying tissue- or condition-specific expression reducing the need for new data collection by repurposing existing large databases.

The system uses a new indexing data structure, which we call Sequence Bloom Tree (SBT), to facilitate searching RNA-seq short-read expression experiments for expressed transcripts. The indexing is independent of the eventual queries, so the approach is not limited to searching only for known genes, but rather can potentially identify arbitrary expressed sequences such as lncRNAs, novel isoforms, or uORFs. The SBT index can be efficiently built, extended, and stored in limited additional space. It also does not require retaining the original sequence files.

Some progress has been made toward enabling sequence search on large databases. The NIH SRA does provide a sequence search functionality [5]; however, this tool requires the selection of a small number of SRA experiments to which to restrict the search and limits the number of reads searched to 2 billion (*≈* 1000 human RNA-seq experiments, or 0.01% of the SRA). Existing full-text indexing data structures such as Burrows-Wheeler transform [4], FM-index [9], compressed suffix arrays [11], wavelet trees [12] or others [18] cannot at present be efficiently built at this scale, and do not directly support the search of millions of short sequences. Word-based indices [1, 17, 25, 26], such as those used by Internet search engines, are not appropriate for edit-distance-based biological sequence search. Sequence-specific solutions caBLAST [15] and caBLASTP [7] have explored using custom compressed indices to speed up search. While successful at protein or genome search, they do not scale to collections of millions of short reads. Further, all of these existing approaches do not handle the additional complication that a match to a query sequence *q* may span many short reads.

The Sequence Bloom Tree can be applied to any set of sequences that can be partitioned into discrete units and allows for fast searching with a greatly reduced storage cost. We apply Sequence Bloom Trees to the problem of searching RNA-seq experiments for expressed isoforms. We build a SBT on 2652 currently publicly available RNA-seq experiments in the SRA for human blood, breast, and brain tissues. Each of these tissues is among the top-5 most common tissues represented in the SRA. Using Sequence Bloom Tree, 2652 short-read sequencing experiments can be searched for a single randomly expressed transcript query in 19 minutes at a reasonable expression threshold. The comparable search time using SRA-BLAST [5] or mapping via STAR [8] is estimated below to be 1.33 days and 1.6 days respectively. We use this newfound speed to search all blood, brain, and breast SRA sequencing runs for all 214,293 known human transcripts. This massive search takes just under 5 days to complete on a single thread.

Sequence Bloom Trees are practical to build: converting from fasta files to sequence bloom filters and building the SBT takes around 2.5 minutes per read set using our mostly single-threaded prototype implementation, a small incremental cost on the generation of sequencing data. They are space efficient as well, requiring less than 4% of the space of the sequences they search. They are accurate, identifying > 77% of truly matching sequencing experiments and with a false positive rate of *<* 21%. They are easily distributed across computers in the cloud or a grid, and they are parallelizable when multiple cores are available. They are also dynamic, allowing insertions and deletions of new experiments, and progressive in the sense that a coarse-grained version of a Sequence Bloom Tree can be downloaded if needed and subsequently refined as more specific results are needed.

We can also view Sequence Bloom Tree as an efficient filter that eliminates many sequencing experiments from consideration to identify the subset of data sets on which more in-depth computational processing can be conducted. For example, a natural downstream processing step would be to run a gene expression estimation algorithm such as Cufflinks [24] or Sailfish [19] on the returned data sets, or to map the reads from the selected experiments using a read mapper such as STAR [8] or Bowtie [13]. The use of Sequence Bloom Tree can drastically reduce the computational effort required for such large-scale processing by ruling out experiments in which the target sequences are not present. By filtering the database before subsequent processing, the subsequent processing time scales relative to the size of the number of hits rather than the size of the database. Accordingly, we compare the running times of Sequence Bloom Tree, SRA-BLAST, and STAR to perform similar filtrations (Section 4.5).

Sequence Bloom Trees have several natural parameters governing the tradeoff between index size, accuracy, and speed. In addition to our computational experiments, we provide some theoretical guidance for setting these parameters to ensure the constructed SBTs perform well.

The main idea behind Sequence Bloom Trees is the creation of a hierarchy of compressed bloom filters, each of which maintains the set of kmers (length-*k* subsequences) present within subsets of the sequencing experiments. The use of a hierarchy of bloom filters is not new: Crainiceanu [6] introduce Bloofi, which maintains a B+-tree of bloom filters of data elements for the application of data management on distributed systems. Specifically, each distributed site has a single filter which stores a set of universally unique identifiers that can be queried from some central B+-tree. Our Sequence Bloom Tree system shares many similarities with Bloofi, particularly the greedy approach to inserting new data elements. However, there are a number of differences as well. First, we apply the idea to the problem of sequence search. Crucially, this allows us to tune the bloom filter error rate much higher than in other contexts (see Theorem 3), vastly reducing the space requirements. We further reduce the space usage (and thus decrease I/O-bound search times) by using bloom filters that are compressed via the RRR [21] compression scheme. RRR bit vectors allow querying a bit without decompression and incur only a *O*(log *m*) factor increase in access time (where *m* is the number of bits in the vector). We also test a batch query system that allows multiple queries to be answered simultaneously.

Sequence Bloom Trees represent a very promising way to index large collections of sequencing experiments to enable fast search by sequence, and here we quantify experimentally their running times, space usage, and accuracy, as well as provide some theoretical guidance for parameter selection. In addition to the application to RNA-seq reported here, we expect SBTs to be useful in the genomic and metagenomic contexts as well. They are our first step toward building a usable, high-performance searchable version of the petabases of sequence data in the SRA.

## 2 Approach

We apply an indexing data structure called Sequence Bloom Tree (Figure 1) to the problem of large-scale sequence search. Sequence Bloom Trees are built around a collection of bloom filters [2, 3]. Bloom filters are an efficient way to store a set of items from a universe 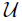. Each filter consists of a bit vector of length *m* and a set of *h* hash functions 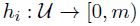 [0, *m*) that map items to bits in the bit vector. Insertion of 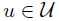 is performed by setting to 1 the bits specified by *h*_*i*_(*u*) for *i* = 1, *… , h*. Querying for membership of *u* checks these same bits; if they are all 1, the filter is reported to contain *u*. Because of overlapping hash results, bloom filters have a one-sided error: they may report an item is present when it is not. This error can typically be made quite small by appropriate choice of *m* and *h*.

**Figure 1:**
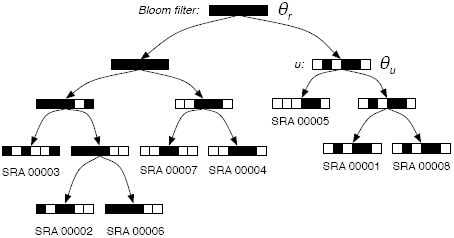
Schematic of a Sequence Bloom Tree. Each node contains a bloom filter containing the kmers present in the sequencing experiments under it.

Sequence Bloom Trees are binary trees in which the sequencing experiments are associated with leaves. Each node *v* of the SBT contains a bloom filter that contains the set of kmers present in any read in any experiment in the subtree rooted at *v*. SBTs are different than cascading bloom filters [23], which aim to reduce false positive rates of a single set query by recursively storing false positives in their own bloom filters.

A primary challenge with scaling Sequence Bloom Trees to terabytes of sequence is saturation of the filters at levels of the tree near the root. The filter at any node *v* is the union of the filters of its children. The union of bloom filters *b*_1_ and *b*_2_ can be efficiently computed by simply taking the bitwise-OR of *b*_1_ and *b*_2_. However, this means as one moves from the leaves to the root, the filters will tend to contain more and more bits set to 1, increasing their false positive rate. This saturation can be overcome using several techniques: appropriate parameter selection (see Section 3.5), grouping of related experiments during insertion into the tree (see Section 3.1), and including only k-mers that have a minimum coverage count (see Section 3.6). Note that filters with poor false positive rates at high levels of the tree only affect query time: accuracy is governed entirely by the false positive rate of the leaf filters.

## 3 Methods

### 3.1 Sequence Bloom Tree construction and insertion

A Sequence Bloom Tree is built by repeated insertion of sequencing experiments. Given a (possibly empty) Sequence Bloom Tree *T*, a new sequencing experiment *s* can be inserted into *T* by first computing the bloom filter *b*(*s*) of the kmers present in *s* and then walking from the root along a path to the leaves and inserting *s* near the bottom of *T* in the following way. When at node *u*, if *u* has a single child, a node representing *s* (and containing *b*(*s*)) is inserted as *u*’s second child. If *u* has two children, *b*(*s*) is compared against the bloom filters *b*(*left*(*u*)) and *b*(*right*(*u*)) of the left *left*(*u*) and right *right*(*u*) children of *u*. The child with the more similar filter under the Hamming distance between the filters becomes the current node, and the process is repeated. If *u* has no children, *u* represents a sequencing experiment *s′*. In this case, a new union node *v* is created as a child of *u*’s parent. This new node has two children: *u* and a new node representing *s*.

As *s* is walked down the tree, the filters at the nodes that are visited are unioned with *b*(*s*). This unioning process can be made fast (and trivially parallelized for large filters) since the union of two bloom filters can be computed by ORing together the bit vectors. This is particularly beneficial where GPU or vector computations can be used for these SIMD operations.

This insertion process is designed to greedily group together sequencing experiments with similar bloom filters. This is important for two reasons. First, it helps to mitigate the problem of filter saturation. If too many dissimilar experiments are present under a node *u*, then *b*(*u*) tends to have many bits set. In addition, by placing similar experiments in similar subtrees, more subtrees are pruned at an earlier stage of a query, reducing query time.

### 3.2 Querying

Given a query sequence *q* and a Sequence Bloom Tree *T*, the sequencing experiments (at the leaves) that contain *q* can be found by breaking *q* into its constituent set of kmers *K*_*q*_ and then flowing these kmers over *T* starting from the root. At each node *u*, the bloom filter *b*(*u*) at that node is queried for each of the kmers in *K*_*q*_. If more than *θ*_*u*_|*K*_*q*_| kmers are reported to be present in *b*(*u*), the search proceeds to all of the children of *u*, where *θ*_*u*_ is a node-specific cutoff between 0 and 1 governing the stringency required of the match. If fewer than that number of kmers are present, the subtree rooted at *u* is not searched further (it is pruned). It has been shown that kmer similarity is highly correlated to the quality of the alignments between sequences [20, 22], and SBT guarantees that if the query sequence is present (at sufficient coverage), it will be found.

When a search proceeds to the children, the children are added to a queue for eventual processing. Even though there may be a large frontier of nodes that are currently active, the memory usage for querying is the trivial amount of memory needed to store the tree topology plus the memory needed to store the single current filter. If querying is parallelized in the natural way by having each thread pull an active frontier node off the single shared queue, the memory usage grows to a single filter per thread, although handling multiple threads simultaneously reading bloom filters from disk needs to be implemented with care to avoid IO contention. The Sequence Bloom Tree timings reported here are all for single-threaded operation.

If several queries are to be made, they can be batched together so that a collection *𝒞* = {*K*_*q*1_, …, *K*_*qt*_} of queries starts at the root, and only queries for which |*b*(*u*) ∩ *K_q_i* | > *θ_u_*|*K_q_i* | are propagated to the children. When 𝒞 becomes empty at a node, the subtree rooted at that node is pruned and not searched further. The main advantage of batching queries in this way is locality of memory references. If *b*(*u*) must be loaded from disk, it need be loaded only once per batch 𝒞 rather than once per query. Batch queries can be parallelized in the same way as non-batched queries by storing with the nodes on the queue the indices of query sets that remain active at that node. Additionally, batch queries offer an alternative means of parallelization where the query collection 𝒞 is split evenly among active threads that merge results for the final query results.

### 3.3 Deletion

Deletion of a leaf from Sequence Bloom Tree proceeds from the to-be-deleted leaf *s* up toward the root. Let *p*(*u*) be the parent of any node *u*, and let *P*_*s*_ be the path from *s* to the root. We can assume that all the nodes along *P*_*s*_ have two children (paths with nodes that have a single child can be compressed into a single node). Assume *s* has a sibling *s′*. To delete *s*, the parent *p*(*s*) is replaced by *s′*, and *p*(*p*(*s*)) has its filter recomputed as the union of *s′* and its other child. We then walk up *P*_*s*_, recomputing each filter as the union of its two child filters. This requires examining only *O*(depth(*s*)) filters. For the experiments reported here, we do not employ deletion.

### 3.4 Theoretical running times for insertion, deletion, and query

Insertion and deletion both follow a single path from the root to a leaf *s*. In each case, two filters per examined node must be read in their entirety to either compute the similarity between the inserted filter and each node’s 2 children, or to recompute the union when deleting. If the filters are of length *m*, the running time for insertion or deletion is *O*(*m* · depth(*s*)). Deletion can be made faster in practice by using lazy deletion: a node is initially only marked deleted and no filters are re-unioned. Once a large enough subtree rooted at *u* has all of its nodes marked as deleted, the filters on *P*_*u*_ are recomputed.

The time for both insertion and deletion depends on the depth of the leaf being added or removed. If insertion and deletion are common, this depth can be reduced by creating a more balanced tree as done in Bloofi [6]. The insertion procedure above favors grouping similar leaves over balance. A modified insertion scheme could favor inserting nodes into smaller subtrees by choosing the child to recurse into based on a combination of filter similarity and subtree size. We do not explore this alternative here as in our use case insertion is rare and deletion never occurs.

### 3.5 Parameter selection

There are two important parameters that need to be set when constructing the bloom filters contained in an Sequence Bloom Tree. These are the bloom filter length (*m*) and the number of hash functions (*h*) used in the filter. We also must set the kmer threshold *θ*_*u*_ for each node. For simplicity here, we use a uniform threshold *θ* = *θ*_*u*_ for all nodes *u*. We explore below the relationship between *m*, *h*, *θ*, and the resulting false positive rate *ξ* of the filters.

#### 3.5.1 Setting the bloom filter false positive rate

Let *S* be a collection of *r* sequencing experiments with the property that each *s* ∈ *S* contains *n* distinct kmers. We first analyze the behavior of a union of filters under the simplifying assumption that the kmer overlap between all pairs of experiments in *S* is uniform. Specifically, assume that the probability that two different experiments *s*_*i*_ and *s*_*j*_ in *S* share any given kmer is *p*. In other words, the expected number of kmers that appear in *s*_*j*_ that do not appear in *s*_*i*_ is *d*(1 *− p*), where *d* is the number of kmers in the experiments. Let *F* be a collection of *r* bloom filters, each of length *m*, where *f*_*s*_ ∈ *F* is the filter containing the kmers from experiment *s* ∈ *S*. Assume each bloom filter employs *h* hash functions. We then can quantify the relationship between the kmer overlap *p* and the saturation of the union filter:

##### Lemma 1.

*Let U* =∪_*f* ∈ *F*_ *f be the union filter of the filters in F as described above. The expected number of items in the union is n* (1 *−* (1 *− p*)^*r*^)*/p.*

*Proof.* We have 𝔼[|*U* |] = 𝔼[|*S*_1_|] + 𝔼[|*S*_2_ *\ S*_1_|] + 𝔼[|*S*_3_ *\ S*_1_ *\ S*_2_|] + *· · ·*. Each kmer in *S*_*i*_ is absent from *∪*_*j*<*i*_*S*_*j*_ independently with probability (1 *− p*)^*i*−*1*^. Therefore 𝔼[|*S_i_ \ ∪_j<i_S_j_*|] = *n*(1 *− p*)^*i*−*1*^, and we have:

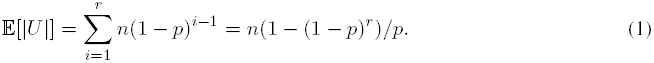

Equation 1 shows, as expected, that when the overlap is large (*p* is close to 1), the number of elements of *U* approaches that of a single filter. Using this relationship, we can select the optimal number of hash functions for such a union as in Theorem 2.

##### Theorem 2.

*The number of hashes that minimizes the FPR of a union filter U with the expected number of elements is*

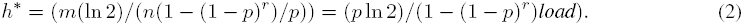

*where load* = *n/m. Under this setting of h, the FPR of U is*

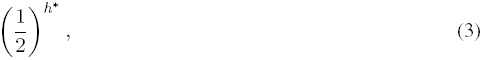

*which is at most* 1/2 *so long as h* ≥* 1.

*Proof.* Follows directly by treating *U* as a single filter containing *n*(1 *−* (1 *− p*)^*r*^)*/p* items.

In the case of Sequence Bloom Trees, we have an advantage that we are not ultimately interested in a single bloom filter query on a kmer, but rather a set of queries of the kmers contained in the longer query string *q*. Thus, we are concerned mostly with the FPR on *queries* rather than FPR on kmers. Theorem 3 explores the connection between the two:

##### Theorem 3.

*Let q be a query string containing ℓ distinct kmers. If we treat the kmers of q as being independent, the probability that >* ⌊*θℓ*⌋ *false positive kmers appear in a filter U with FPR ξ is*

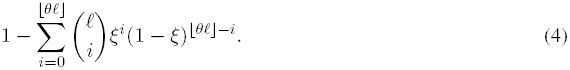

*The above expression is nearly* 0 *when ξ* ≪ *θ.*

*Proof.* Treating each kmer in *q* independently allows us to model the repeated queries using a binomial distribution, yielding (4). A false positive in *q* occurs when > ⌊*θℓ*⌋ false positive kmers occur in *U*. Let *X* be the number of false positive kmers, and let *Y* be the number of correctly determined kmers. Then Pr[*X > θℓ*] = Pr[*Y ≤ ℓ − θℓ*]. When *θ ≥ ξ*, we have *ℓ − θℓ ≤ ℓ*(1 *− ξ*) = *E*[*Y* ], and the following bound holds by Chernoff’s inequality:

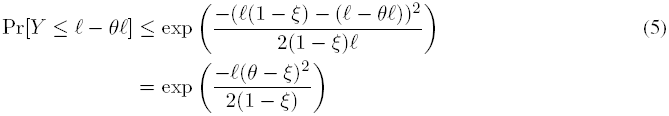

In our search application it is natural to require that at least 1/2 the kmers of a query are present; if < 1/2 are present it is fair to say that the query is not contained within the experiment. In this case, the threshold for the fraction of kmers that must be present, *θ*, is *≥* 0.5, and so if we choose the FPR of the bloom filters to be 0.5, by Theorem 3, we will be unlikely to observe *> θ* fraction of false positive kmers in the filter. A bloom filter FPR of 0.5 is much higher than typical applications of bloom filters, in which very low false positive rates are sought. The above analysis assume independence of the kmers, which is of course unrealistic. Nevertheless, it formalizes the intuition that choosing a high FPR leads to few errors. By choosing such a high filter FPR, we can use smaller filters, limiting the memory footprint of the Sequence Bloom Tree.

#### 3.5.2 A sampling scheme to avoid saturation

The case when *h* <* 1 in Theorem 2 commonly occurs in practice when applying Sequence Bloom Trees because either the load of each filter is high or the overlap *p* is low. While using fewer than 1 hash function is not possible, we can simulate having *h* <* 1 effective hash functions by randomly sampling only a *h\** fraction of the kmers that would be added to *U* (i.e. by adding a kmer to *U* with probability 0 *< h* <* 1). This is possible here because when searching for a query string *q*, we require many kmers of *q* to be present in the filter. The effect of omitting kmers due to random sampling therefore can be mitigated (up to a point) by lowering the threshold *θ* of fraction of kmers that are required to be present for the query to pass the filter. In this situation, the bloom filter can be regarded has having two-sided error: its FPR is *f* as described above and its false negative rate is now (1 *− h\**)(1 *− ξ*), since a false negative occurs if the kmer was not added (which occurs with probability (1 *− h\**)) and this true negative query (from the perspective of the filter) was not reported as present due to a false positive (which occurs with probability (1 *− ξ*)). Note here that a high false positive rate works to our benefit in reducing false negatives due to sampling. This leads to the following corollary:

##### Corollary 4.

*If kmers are sampled with rate h* <* 1, *using cutoff θ′* = (1 *−* (1 *− h\**)(1 *− ξ*))*θ will, with high probability, achieve the same number of false positive queries as using θ with no sampling.*

*Proof.* If *θℓ* kmers of the query are truly present in the subtree, then we expect at least 1 *−* (1 *− h\**)(1 *− ξ*))*θℓ* to be present after sampling.

For the experiments reported here, we do not perform this sampling, as it is not needed. For even larger collections of sequencing experiments, however, this sampling scheme may prove useful.

### 3.6 Building bloom filters

In the experiments here, bloom filters were constructed using the Jellyfish kmer counting library [16] from short-read FASTA files downloaded from the NIH SRA by counting canonical kmers (the lexicographically smaller kmer between a kmer and its reverse complement). We choose *k* = 20 as these kmers are reasonably unique within the human genome.

To avoid counting kmers that result from sequencing errors and to attempt to select kmers from genes with reasonable coverage, we built trees containing kmers that occur greater than a file-dependent threshold. This threshold *count*(*s*_*i*_) was determined using the file size of experiment *s*_*i*_ as follows: *count*(*s*_*i*_) = 1 if *s*_*i*_ is 300 MB or less, *count*(*s*_*i*_) = 3 for files of size 300–500 MB, *count*(*s*_*i*_) = 10 for files of size 500 MB–1 GB, *count*(*s*_*i*_) = 20 for files between 1 GB and 3 GB, and *count*(*s*_*i*_) = 50 for files > 3 GB or larger FASTA files. These cutoffs were determined via the analysis of a small set of 18 sequence experiments of various sizes and tissue types and were chosen such that at least 60% of the transcripts expressed at a non-zero level in each of these files had an estimated uniform coverage above this number. While this is not a statistically significant sample relative to the size of the database, in practice we found these numbers to outperform two naïve thresholds (*count*(*s*_*i*_) = 0 and *count*(*s*_*i*_) = 3 for all *i*) in speed and accuracy. We report only the results from the file-dependent threshold for this reason.

We can use a cutoff based on file size here because all the experiments sequenced the human transcriptome. In a situation where experiments of mixed organism origin are included, a more sophisticated scheme based directly on sequencing coverage would be needed to avoid counting sequencing errors.

We use a filter FPR of 0.5 and *h* = 1 as suggested by the above theorems. This leads to an optimal, uncompressed filter size of 239 MB, and any kmer of sufficient coverage that is shared between two files will correspond to a shared bit. After the Sequence Bloom Tree is built, the filters (both leaf and internal) are compressed using the RRR [21] bit vector compression scheme as implemented in the SDSL [10].

### 3.7 Representative query sets and ground truth results

To determine the accuracy of Sequence Bloom Trees, we selected a subset of 100 random sequence files and used Salmon (the latest version of Sailfish [19]) to quantify the expression of all transcripts in each of these experiments. All Sequence Bloom Tree queries are queried on the full set of 2652 files but the accuracy below is computed based only on the random subset of files for which we computed expression results from Salmon.

Three collections of representative queries were constructed, denoted by *High*, *Medium*, and *Low*, that include transcripts that are likely to be expressed at a higher, medium, or low level in at least one experiment contained in the tree. These collections were created using the gene expression information from the ground truth subset run through Salmon. Specifically, the *High* set was chosen to be 100 random transcripts of length > 1000 nt with an estimated abundance of > 1000 transcripts per million (TPM). The *Medium* and *Low* query sets were similarly chosen randomly from among transcripts with > 500 and > 100 TPM, respectively. These Salmon estimates were taken as the approximate ground truth of expression for the query transcripts.

### 3.8 SRA-BLAST

There are presently no search or alignment tools that can solve the sequence search problem in short-read sequencing files at the scale we attempt here. However, as alignments can be used to determine query coverage and thus the presence of transcripts in sequence files, we compare with SRA-BLAST [5]. SRA-BLAST has a strict limitation on the total nucleotide count that can be searched at once and requires specifying SRX (experiment) files rather then SRR (run) files. Because it is impractical to use SRA-BLAST at the SBT scale, we estimated an average SRA-BLAST query time from twenty random queries of a transcript against a single SRX experiment set using the SRA-Blast webtool [5]. Specifically we randomly selected a short read file from the total 2652 set and a query from the *Low* representative query set and recorded the time it took SRA-BLAST to return an alignment. As the representative query set does not guarantee expression in the full 2652 set, this provides a more reasonable estimate that accounts for both trivial misses and more expensive hits. Some publicly available SRR files cannot be searched using this webtool and random queries containing these files were discarded. The extrapolated time to process one > 1000 nt query against one megabase of sequence read file is recorded at 0.0144 (seconds per-megabase-per-query).

### 3.9 STAR

We also compare our search times with an alignment-based approach using a read mapping algorithm, STAR [8]. To do this, we constructed a STAR index each set of 100 sequence queries and mapped the reads in each of the 100 files analyzed by Salmon (our ground truth set) to the queries allowing 0 mismatches (unsorted BAM output, size 11 pre-index string). An average STAR query time of 0.0013 seconds per-megabase-per-query was calculated from these results by normalizing the time it takes to perform a STAR alignment against the total size of each sequence file and number of queries. In all queries, zero alignment mismatches were allowed and STAR was allowed to use 15 parallel threads. Note that the time per-megabase-per-query does not include the time to build the STAR index and that STAR was allowed to process all 100 queries simultaneously in a batch while the SBT and SRA-BLAST were not. A separate analysis was run to compare the batch timings for SBT.

## 4 Results

### 4.1 Construction of a large Sequence Bloom Tree containing human RNA-seq experiments

A Sequence Bloom Tree was constructed from 2652 human, RNA-seq short-read sequencing runs from the NIH SRA. These 2652 files represent the entire set of publicly available, human RNA-seq runs from blood, brain, and breast tissues stored at the SRA as determined by keywords in their metadata and excluding files sequenced using the SOLID technology. Files where the metadata was unclear about tissue type or experimental setup were discarded. This tree was used for all experiments described below except the all transcript analysis which was run on a slightly larger 2744 file set.

All times in these experiments were obtained on a shared computer with 16 Intel Xeon 2.60GHz CPUs using a single thread (or 15 threads in the case of STAR). SBT queries were limited to the size of a single compressed filter at any one time which was strictly less than 239 MB (though usually much less in practice).

### 4.2 Time and space to construct and store the bloom tree

The construction of the Sequence Bloom Tree involves 3 major tasks: creation of bloom filters for each of the experiments included at its leaves, the construction of the tree and internal bloom filters, and the RRR compression of each of the filters. The leaf filter construction takes the longest time taking just under 3 days using Jellyfish’s counting library [16]. Once the leaf filters have been constructed, the tree can be built in under 20 hours using a single thread. The filters in the tree can be compressed in an additional 14 hours, and once built can be searched and stored without decompression. As the entire build process need only be done once (and additional insertions / deletions can be performed to modify an existing tree without rebuilding), these times are reasonable for the scale of data being processed. Since even with a single thread, this construction time is only approximately 2.5 minutes per read file, the time to build the index is a small marginal cost when creating a new sequencing file.

The SBT is designed to be efficient in both space and time. The complete set of sequence reads searched here was originally 5 TB in size. The equivalent compressed bloom filters at the leaves of the corresponding Sequence Bloom Tree are only 63 GB in size. Because the entire Sequence Bloom Tree can be reconstructed from these leaves by repeated insertion, the Sequence Bloom Tree can be transferred between (say) a sequence database and a user using only this 63 GB. The entire Sequence Bloom Tree, including internal filters, requires only 200 GB, a small fraction of the size of the original sequencing data. Table 1 gives the compressed and uncompressed sizes of the raw sequence files, the bloom tree, and the leaves. SBT is not meant to replace existing database storage techniques but merely offers a much more efficient means of querying large files without downloading the full dataset.

**Table 1:**
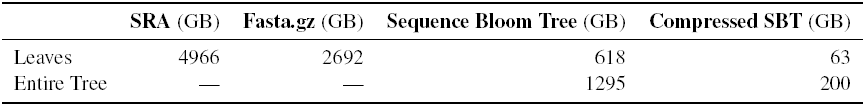
Storage costs by file type. Leaves gives the size of the data associated with the leaves; the SRA and Fasta.gz columns give the storage cost for the 2652 input sequence files (original download and standard storage) while the others gives the storage cost of the 2652 leaf filters. Entire Tree gives the total storage cost for the leaves and all internal nodes.

|  | SRA (GB) | Fasta.gz (GB) | Sequence Bloom Tree (GB) | Compressed SBT (GB) |
| --- | --- | --- | --- | --- |
| Leaves | 4966 | 2692 | 618 | 63 |
| Entire Tree | — | — | 1295 | 200 |

### 4.3 Query speed

SBT was designed to permit efficient sequence-based querying of collections of genomic experiments. To explore the parameter space of the Sequence Bloom Tree, each of the *High*, *Medium*, and *Low* sets defined above were queried in the Sequence Bloom Tree as batch queries (providing all 100 queries at once) for various kmer thresholds *θ* = 0.5, 0.6, 0.7, 0.8, 0.9, 1.0. In all case, SBT is able to process these queries in under 25 seconds per query using a single processing thread and with a memory footprint less than the size for a single uncompressed filter (239 MB). The results are shown in Figure 2. As the running time is robust to different TPM queries and the total query time is relatively the same regardless of kmer threshold, we select an optimal threshold of 0.8 and an optimal minimum query TPM of *Low* based on its ratio of true positive to false positive (Section 4.4).

**Figure 2:**
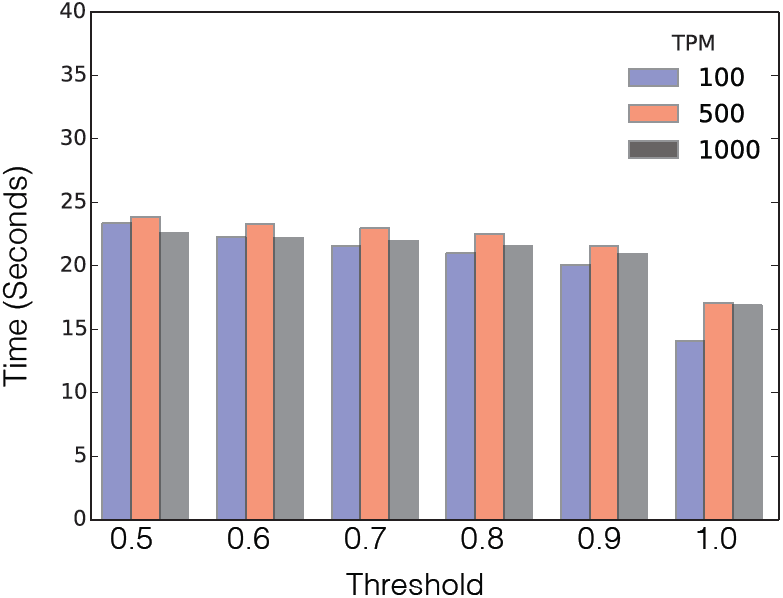
Average time to query a > 1000 nt sequence across 2652 SRA files indexed in a Sequence Bloom Tree (using 100 query batches). The query threshold *θ* for a hit varies from 0.5 to 1 without a significant change in processing time and time is largely independent of TPM.

Using these parameter values, we ran the *Low* queries individually to determine an average user time to find all matches within the database without the advantage of batched queries and compare our time to similar searches run through SRA-BLAST and STAR (Figure 3A). Because SRA-BLAST lacks an automated querying mechanism, this time is estimated by extrapolating the time to perform 20 queries to get an estimated time per megabase of searched sequence (Section 3.8). Likewise a STAR index was created for the query set (e.g. *Low*), and each of the sequence read files was aligned to these queries using STAR (15 threads). Because this is impractical for 2652 files, we extrapolate the estimated running time from a set of 100 randomly chosen files. The reported running times for STAR are always batched and exclude the creation of the index, but unlike with Sequence Bloom Tree, the index for STAR needs to be rebuilt for each new query set. Despite these handicaps, the SBT is still roughly 100x faster than competing metrics using one thread and over 10x faster than the 15-thread run.

**Figure 3:**
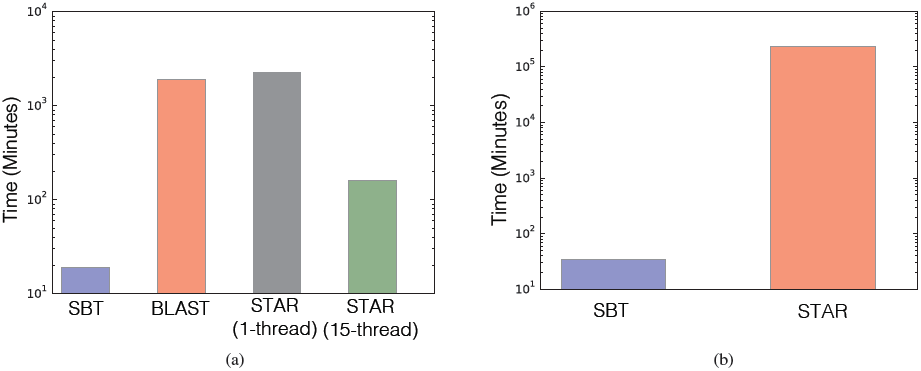
**(A)** Running times of search tools for one genomic query. The Sequence Bloom Tree time is the per-query time to search over the SBT with 2652 leaves using a maximum of a single filter in active memory and one thread. The other bars show the estimated time to achieve the same query results using SRA-BLAST- and mapping-based approaches. SRA-BLAST was run on the publically available web-tool and STAR was allowed to run on both 1 and 15 threads. **(B)** Running times of search tools for 100 batched genomic queries. In both cases SBT and STAR were run using one thread and SBT was limited to a single filter in RAM.

When we allow SBT to batch the same query set as STAR the speed advantage of Sequence Bloom Tree becomes even more apparent (Figure 3B). As the majority of the SBT processing time is devoted to the I/O of loading and unloading bit vectors from memory, the batch approach which amortizes this cost over 100 simultaneous lookups is over 6715*×* faster than the corresponding STAR. As SRA-BLAST was not run locally, it was not compared in a batched fashion.

While no previous system solved the large-scale, short-read search problem that we consider here, the natural alternatives tested above (SRA-BLAST and a fast read mapper) are impractical for searching collections of sequences of the size that Sequence Bloom Tree can handle.

### 4.4 Query accuracy

The error in the bloom filters is one sided, so if a kmer is present within the reads with a high enough count, the bloom tree will correctly report its existence. However, this does not necessarily imply that all queries that are expressed in the experiment are correctly found. This is because the bloom tree query requires that at least *θ* fraction of the kmers are present at a sufficient count. For example, a transcript with high read coverage across 75% of its length will only be found by the Sequence Bloom Tree at *θ* ≤ 0.75 while expression quantification estimators such as Sailfish may be more relaxed. This results in a discrepancy between the “expressed” transcripts that an expression quantification estimator such as Sailfish finds and what the Sequence Bloom Tree can see, leading to some false negatives. On the other hand, false positives can also arise due to bloom filter errors or kmers that are present in several transcripts.

To analyze the accuracy of the Sequence Bloom Tree filter, we quantify true positive and false positive rates over the subset of 100 randomly selected leaves used to generate the *High*, *Medium*, and *Low* query sets. Ideally, we would assess this accuracy over the entire leaf set, but it is at present impractical to perform transcript quantification over 2652 short-read sets. In all cases, we query the entire bloom tree but build our accuracy results only from the subset of files passed through the Salmon quantification tool. Figure 4 shows ROC curves for the three query sets as *θ* is varied. In this plot, a true positive (TP) is a leaf for which Salmon reports that the query is expressed and the Sequence Bloom Tree returns this leaf in its results. A false positive (FP) is a leaf for which Sequence Bloom Tree returns the leaf, but Salmon does not report that the query transcript is expressed at a sufficiently high level (TPM ≥ 100, 500, 1000 for *Low*, *Medium*, *High*, respectively). The curves are the average over the 100 queries within a query set. With a FP rate of 0.17 on average nearly 80% of the true results are found by Sequence Bloom Tree with an optimal ratio of 0.8. The performance on the various query sets is comparable.

**Figure 4:**
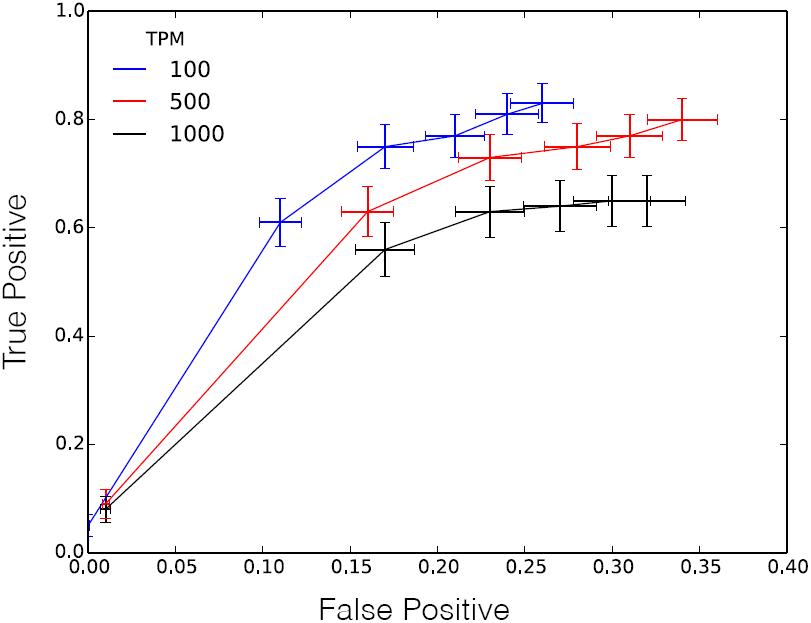
Receiver operating characteristic curve averaged over 100 queries from the *High*, *Medium*, and *Low* query sets. The ROC curve was created by repeatedly querying the same randomly selected query set using different thresholds *θ* for total k-mer coverage from 0.5 to 1.0. Crosses denote the standard error about the mean for both axes.

While these accuracy results represent a good tradeoff given the vastly increased speed of Sequence Bloom Tree, the results are in fact better than shown by Figure 4 because there are a few outlier queries for which the Sequence Bloom Tree reports no results but their expression is above the TPM threshold as estimated by Salmon. Figure 5 makes this clear: it plots the median TP and FP rates (rather than the average) for each query set. For most queries, 100% of the correct experiments are found. The area under the median ROC curve is 0.97, 0.95 and 0.94 for the three tested TPM levels respectively. For a minority of the experiments, no correct queries are returned. This is a result of the threshold used to count kmers within each file: if the kmer coverage of a transcript in an experiment *s*_*i*_ is less than *count*(*s*_*i*_), Sequence Bloom Tree will not find it, while Salmon can still give it a high relative expression estimation.

**Figure 5:**
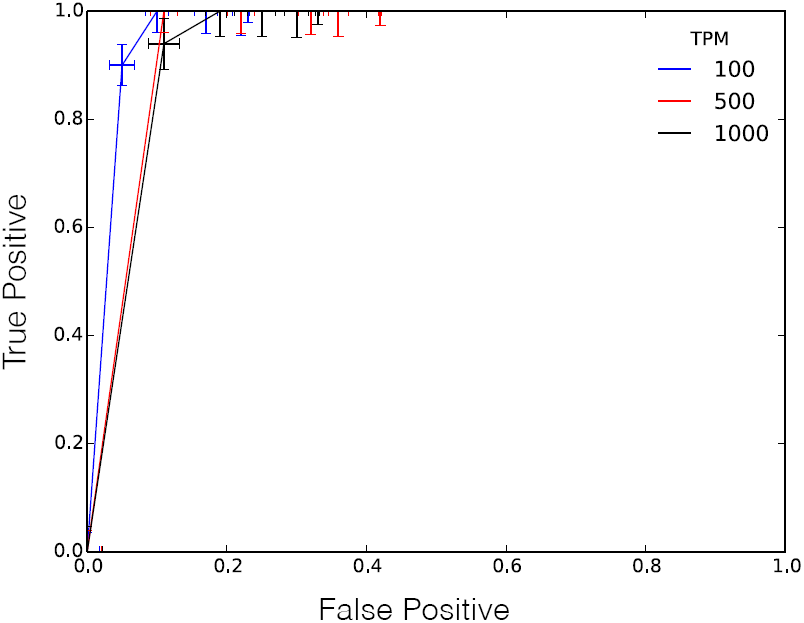
ROC curves for the median (rather than average as in Figure 4) over 100 queries from the *High*, *Medium*, and *Low* query sets as the threshold *θ* is varied from 0.5 to 1.0. Crosses denote the standard error about the mean for both axes

An experimentally derived optimal *θ* of 0.8 was determined by maximizing the ratio between true positive and false positive rates. At this setting and for > 1000 nt queries with TPM above 100, a true positive rate of 0.8 is achieved against a false positive rate of under 0.2. The returned leaf set includes almost all of the true query hits while adding only a small fraction of the overall search space as false positives. If the Sequence Bloom Tree is used as a screen for a subsequent search or alignment tool, the only side effect from a false positive is an additional cost to the query time proportional to the number of files mistakenly searched (see Section 4.5).

These results, combined with the timing results of the previous section, show that large collections of short-read files can be searched for transcript expression with high accuracy quickly, a task that was not previously possible.

### 4.5 Additional benefit as an alignment filter

We can also view the SBT as a filter that eliminates files that do not contain the query with high confidence. By running SBT before an alignment tool on the remaining set, the total search space can be reduced without a significant loss of accuracy. To demonstrate this, we estimate the time it would take STAR or SRA-BLAST to process the hits found by SBT rather then the full database (Figure 6). These values were extrapolated from the relative search space trimmed from the random set (35% of the original database size using the *Low* query set) and represent a 3× speedup over the naïve approach. As less than 20% of this remaining data represents false positives, the SBT filter the database on the scale of the number of hits rather than the total database size. These results represent some of the worst case behaviors of the SBT. If query hits are rare or the SBT is built over a larger range of tissue types with distinct expression patterns, a greater number of files will be filtered by SBT and the speedup will grow accordingly.

**Figure 6:**
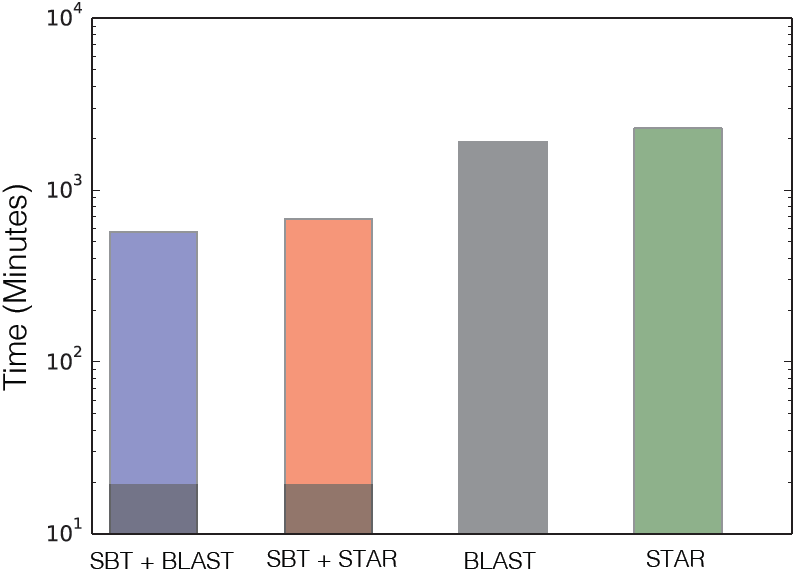
Estimated time to process the full 2652 dataset using a combination of SBT (as a filter) followed by STAR or SRA-BLAST versus the time to process STAR or SRA-BLAST naïvely. Dark bars denote the SBT time while the total height represents the total time using conventional search tools over the remaining set.

### 4.6 The benefit of the tree structure

To assess the utility of the tree of internal bloom filters in the Sequence Bloom Tree, we plot the number of nodes of the tree visited versus the number of leaves in which the query was found (Figure 7). Naturally, when a query is found in many of the leaves, the query must also visit a nearly equal number of internal tree nodes, and so the tree structure would not provide any benefit over merely searching all the leaf filters directly. On the other hand, when the query is found in only a few leaves, the total number of nodes visited can be significantly smaller than the number of leaves. For the SBT built here, we find that for queries that are found in 600 or fewer leaves, the tree structure and internal nodes result in an improvement of overall efficiency by visiting fewer than 2652 nodes. We do find that a sizable minority of human transcripts are found in > 600 leaves (section 4.7), however, it is not possible to identify those queries ahead of time to avoid searching the internal SBT nodes for them alone.

**Figure 7:**
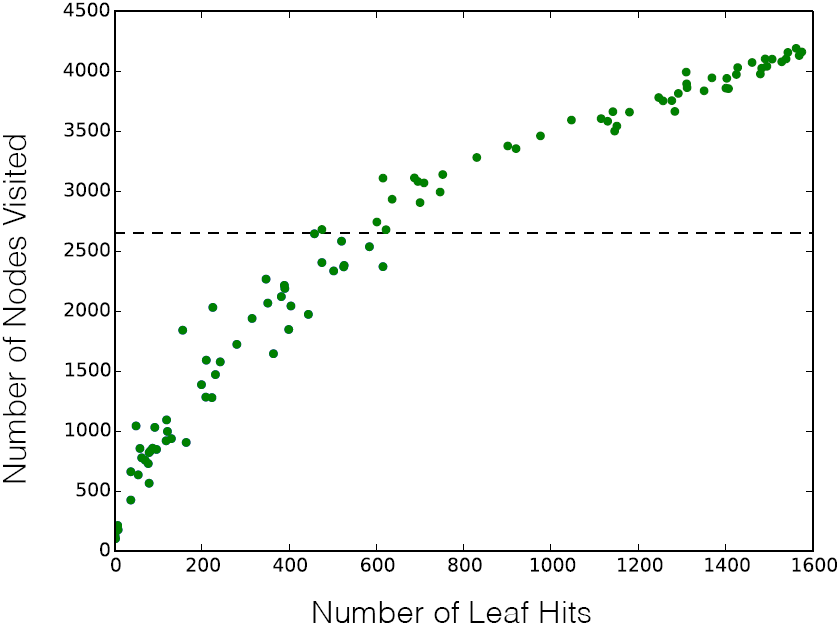
Total number of Sequence Bloom Tree nodes visited as a function of the number of leaf hits. A naïve approach that did not use the tree would require querying 2652 leaf filters for all queries (denoted by dashed line). Over half of the randomly selected queries fall below this threshold.

### 4.7 An enormous query set

To demonstrate the large-scale analysis that can be conducted with Sequence Bloom Tree, we queried the entire set of 214,293 known human transcripts against a larger 2744 file Sequence Bloom Tree. This is a scale of analysis that was not previously possible, and it shows the potential for uncovering expression patterns of transcripts in an unbiased way. The timing information for these queries can be found in Figure 8. At *θ* = 0.7 this can be completed using a single thread in just under 5 days. In contrast, running Salmon (single-threaded) on all these files would take an estimated 81.6 days.

**Figure 8:**
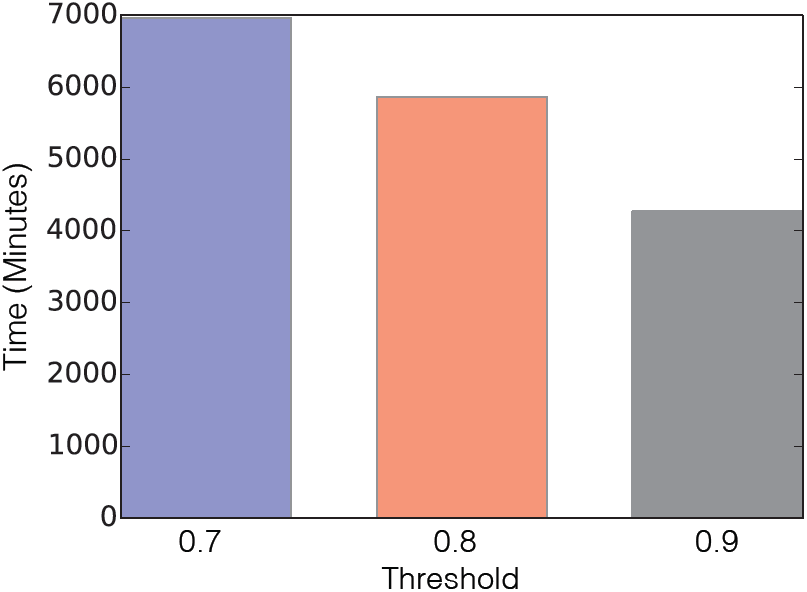
Total times (single-threaded) for querying all 214,293 human transcripts against all publicly available blood, breast, and brain RNA-seq experiments in the SRA for *θ* = 0.7, 0.8, 0.9

Using this, we can get an estimate for the distribution of how widely expressed the set of human transcripts are. That distribution is shown in Figure 9. Even at the most relaxed threshold tested (*θ* = 0.7), the maximum hit count for a single query was 2321 out of 2744 files. We can also estimate the number of tissue specific genes expressed in our dataset. We define a tissue-specific transcript to be one in which over 70% of the matching leaves belonged to a single tissue and, conversely, the transcript was also found in over 33% of the leaves for that tissue. Table 2 gives the number of such blood-, brain-, and breast-specific transcripts.

**Table 2:**
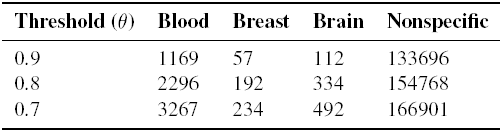
Tissue-Specific Expression in Human Transcriptome. Transcripts were considered tissue specific if they were found in > 33% of the sequencing runs for a tissue and additionally ≥70% of the runs in which the transcript was found were that tissue.

| Threshold ($\theta$) | Blood | Breast | Brain | Nonspecific |
| --- | --- | --- | --- | --- |
| 0.9 | 1169 | 57 | 112 | 133696 |
| 0.8 | 2296 | 192 | 334 | 154768 |
| 0.7 | 3267 | 234 | 492 | 166901 |

**Figure 9:**
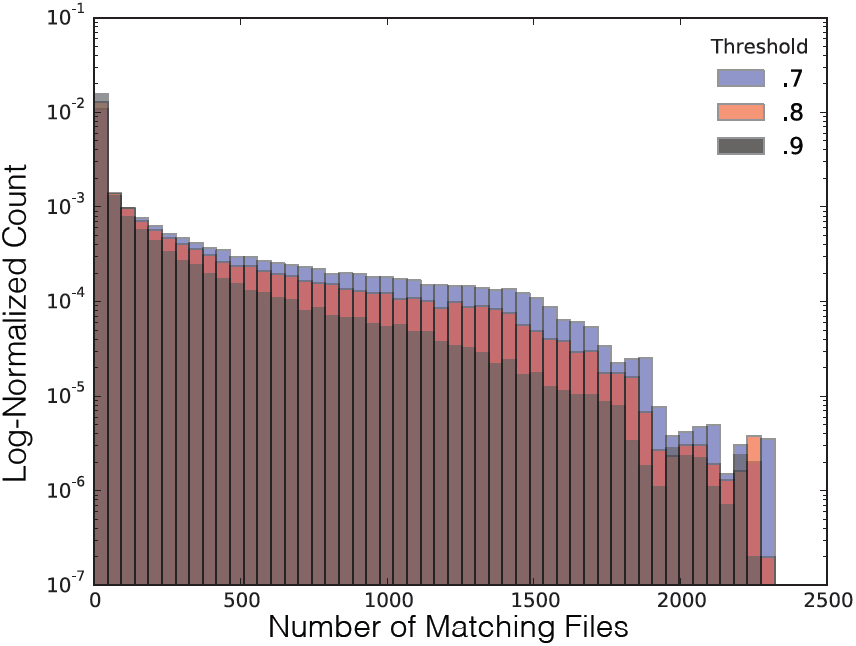
Distribution of hit counts for each query as a function of threshold.

## 5 Conclusion

The Sequence Bloom Tree represents a novel approach to sequence searching that processes data several orders of magnitude faster then existing methodologies. While the SBT can be used for a variety of genomic datasets, we demonstrate the effectiveness of the method as an accurate estimator of transcript presence and as a screening process for subsequent analyses. Our results show that the Sequence Bloom Tree is efficient in terms of time and space, drastically outperforming other existing natural approaches for searching large collections of short reads.

As noted in Crainiceanu [6], the problem of bloom filter saturation — where most of the filter bits are set to 1 leading to a large filter FP error rate — is the primary challenge with scaling hierarchies of bloom filters to larger data sets. We tackle this challenge here by exploiting the fact that each query string consists of many kmers that can be independently queried in the bloom filter and so the bloom filter error probability can be increased and hence its size decreased.

We also combat saturation by only counting kmers that occur a sufficient number of times as determined by the *count*(*s*_*i*_) function. This is a reasonable approach when searching for expressed genes among high-coverage RNA-seq experiments. However, this limits the technique’s ability to find lowly expressed genes, and the thresholds chosen for *count*(*s*_*i*_) introduce an arbitrary distinction between high- and low-expressed genes. Removing or improving this limitation is an important direction for future work.

An open-source prototype implementation of Sequence Bloom Tree is available at http://www.cs.cmu.edu/∼ckingsf/software/bloomtree.

## Acknowledgement

We would like to thank Darya Filippova, Hao Wang, Emre Sefer, Greg Johnson, and especially Guillaume Marçais, Geet Duggal, and Rob Patro for sharing their source code, for valuable discussions, and comments on the manuscript.

**Funding:** This research is funded in part by the Gordon and Betty Moore Foundation’s Data-Driven Discovery Initiative through Grant GBMF4554 to Carl Kingsford. It is partially funded by the US National Science Foundation (CCF-1256087, CCF-1319998) and the US National Institutes of Health (R21HG006913, R01HG007104). C.K. received support as an Alfred P. Sloan Research Fellow.

## References

1 R. Baeza-Yates and B. Ribeiro. Modern Information Retrieval. Addison-Wesley, 1999.

2 Burton H. Bloom. Space/time trade-offs in hash coding with allowable errors. Communications of the ACM, 13(7):422–426, 1970.

3 Andrei Broder and Michael Mitzenmacher. Network applications of bloom filters: A survey. Internet Mathematics, 1(4):485–509, 2005.

4 Michael Burrows and David J. Wheeler. A block sorting lossless data compression algorithm. Technical Report 124, Digital Equipment Corporation, 1994.

5 Christiam Camacho, George Coulouris, Vahram Avagyan, Ning Ma, Jason Papadopoulos, Kevin Bealer, and Thomas L Madden. Blast+: architecture and applications. BMC Bioinformatics, 10(1): 421, 2009.

6 Adina Crainiceanu. Bloofi: a hierarchical bloom filter index with applications to distributed data provenance. In Proceedings of the 2nd International Workshop on Cloud Intelligence, page 4. ACM, 2013.

7 Noah M. Daniels, Andrew Gallant, Jian Peng, Lenore J. Cowen, Michael Baym, and Bonnie Berger. Compressive genomics for protein databases. Bioinformatics, 29(13):i283–i290, 2013. doi: 10.1093/bioinformatics/btt214.

8 Alexander Dobin, Carrie A. Davis, Felix Schlesinger, Jorg Drenkow, Chris Zaleski, Sonali Jha, Philippe Batut, Mark Chaisson, and Thomas R. Gingeras. STAR: ultrafast universal RNA-seq aligner. Bioinformatics, 29(1):15–21, 2013. doi: 10.1093/bioinformatics/bts635.

9 Paolo Ferragina and Giovanni Manzini. Indexing compressed text. Journal of the ACM, 52(4): 552–581, 2005.

10 Simon Gog, Timo Beller, Alistair Moffat, and Matthias Petri. From theory to practice: Plug and play with succinct data structures. In 13th International Symposium on Experimental Algorithms (SEA 2014), pages 326–337, 2014.

11 Roberto Grossi and Jeffrey Scott Vitter. Compressed suffix arrays and suffix trees with applications to text indexing and string matching. SIAM Journal on Computing, 35(2):378–407, 2005.

12 Roberto Grossi, Jeffrey Scott Vitter, and Bojian Xu. Wavelet trees: From theory to practice. In Data Compression, Communications and Processing (CCP), 2011 First International Conference on, pages 210-221. IEEE, 2011.

13 B Langmead, C Trapnell, M Pop, and SL Salzberg. Ultrafast and memory-efficient alignment of short DNA sequences to the human genome. Genome Biol, 10:R25, 2009.

14 Rasko Leinonen, Hideaki Sugawara, Martin Shumway, and The International Nucleotide Sequence Database Collaboration. The sequence read archive. Nucleic Acids Res., 39(Database issue): D19–D21, 2011.

15 P-R Loh, M Baym, and B Berger. Compressive genomics. Nature Biotechnology, 30(7):627–630, 2012.

16 Guillaume Marçais and Carl Kingsford. A fast, lock-free approach for efficient parallel counting of occurrences of k-mers. Bioinformatics, 27(6):764–770, 2011. doi: 10.1093/bioinformatics/btr011.

17 G. Navarro, E. Moura, M. Neubert, N. Ziviani, and R. Baeza-Yates. Adding compression to block addressing inverted indexes. Information Retrieval, 3:49–77, 2000.

18 Gonzalo Navarro and Veli Mäkinen. Compressed full-text indexes. ACM Comput. Surv., 39(1), April 2007. ISSN 0360-0300. doi: 10.1145/1216370.1216372.

19 R. Patro, S. M. Mount, and Carl Kingsford. Sailfish enables alignment-free isoform quantification from RNA-seq reads using lightweight algorithms. Nature Biotechnology, 32:462–464, 2014.

20 Nicolas Philippe, Mikaël Salson, Thérèse Commes, and Eric Rivals. CRAC: an integrated approach to the analysis of RNA-seq reads. Genome Biology, 14(3):R30, 2013.

21 Rajeev Raman, Venkatesh Raman, and S. Srinivasa Rao. Succinct indexable dictionaries with applications to encoding k-ary trees and multisets. In Proceedings of the Thirteenth Annual ACM-SIAM Symposium on Discrete Algorithms, SODA ’02, pages 233–242, Philadelphia, PA, USA, 2002. Society for Industrial and Applied Mathematics.

22 Kim Rasmussen, Jens Stoye, and Eugene Myers. Efficient q-gram filters for finding all E-Matches over a given length. Journal of Computational Biology, 13(2):296–308, 2006.

23 Kamil Salikhov, Gustavo Sacomoto, and Gregory Kucherov. Using cascading Bloom filters to improve the memory usage for de Brujin graphs. Algorithms for Molecular Biology, 9:2, 2014.

24 Cole Trapnell, Adam Roberts, Loyal Goff, Geo Pertea, Daehwan Kim, David R Kelley, Harold Pimentel, Steven L Salzberg, John L Rinn, and Lior Pachter. Differential gene and transcript expression analysis of RNA-seq experiments with TopHat and Cufflinks. Nat. Protocols, 7(3): 562-578, 03 2012.

25 I. Witten, A. Moffat, and T. Bell. Managing Gigabytes. Morgan Kaufmann Publishers, 2nd edition, 1999.

26 N. Ziviani, E. Moura, G. Navarro, and R. Baeza-Yates. Compression: A key for next-generation text retrieval systems. IEEE Computer, 33:37–44, 2000.

